# Analysis of Transcripts and splice isoforms in Red Clover (*Trifolium pratense* L.) by single-molecule long-read sequencing

**DOI:** 10.1101/330977

**Authors:** Yuehui Chao, Jianbo Yuan, Sifeng Li, Siqiao Jia, Liebao Han, Lixin Xu

**Author notes:** Corresponding author: Turfgrass Research Institute, College of Forestry, Beijing Forestry University, Beijing 100083, China. E-mail: Liebao Han; Lixin Xu.

## Abstract

Red clover (*Trifolium pratense* L.) is an important cool-season legume plant, which is the most widely planted forage legume after alfalfa. Although a draft genome sequence was published already, the sequences and completed structure of mRNA transcripts remain unclear, which limit further explore on red clover. In this study, the red clover transcriptome was sequenced using single-molecule long-read sequencing to identify full-length splice isoforms, and 29,730 novel isoforms from known genes and 2,194 novel isoforms from novel genes were identified. A total of 5,492 alternative splicing events was identified and the majority of alter spliced events in red clover was corrected as intron retention. In addition, of the 15,229 genes detected by SMRT, 8,719 including 1,86,517 transcripts have at least one poly(A) site. Furthermore, we identified 4,333 long non-coding RNAs and 3,762 fusion transcripts. Our results show the feasibility of deep sequencing full-length RNA from red clover transcriptome on a single-molecule level.

## Introduction

Red clover (*Trifolium pratense* L.) is an important legume plant that plays a key role in sustainable intensification of livestock farming systems (1). Red clover is native to Europe, Western Asia and northwest Africa, but it is planted in many other regions because it is adapted to a wide range of soils (2, 3). As a legume plant, it has a higher protein content because of nitrogen fixation by nodulation via symbiosis with the soil microbe *Rhizobium leguminarosum*, and its reduced need for nitrogen fertilizer input can reduce the environmental footprint of grassland-based agriculture (4). Red clover is a diploid (2n = 14) species that is naturally cross-pollinated, with a genome assembled with 309 Mb in 39,904 scaffolds reported recently (5). Next-generation high-throughput sequencing (NGS), known as transcriptome analysis or RNA-Seq, has emerged as a revolutionary tool to better understand differential gene expression and regulatory mechanisms. In this approach, there is no strict requirement for a reference genome sequence, so it is suitable for model or non-model species. With this tool, works on transcriptome analysis were accomplished in red clover. In 2014, RNA-Seq and analysis of red clover leaves of drought and non-drought plants provided a rich source for gene identification and the genetic basis of drought tolerance (6). A *de novo* transcriptome assembly of red clover was used for genome-wide identification of different plant transcription factor families, gene expression analysis of different tissues and dynamic spatial gene coexpression networks (7). As a powerful tool for description of gene expression levels and individual splice junctions, short-read RNA sequencing has been becoming focus of scientific researchers, but this tool cannot provide full-length sequence and alternatively spliced forms for each RNA. For example, in *Arabidopsis thaliana*, alternatively spliced forms exist in more than 80% multiple-exon genes (8). Information about RNA sequence and alternative spliced forms is crucial for deeply understanding plant transcriptome and their potential biological consequences. Pacific Biosciences single molecule long reads sequencing technology (SMRT), also called the 3^rd^ generation sequencing technology, is applied to effectively capture full-length sequence of genome and transcripts and accurately identify full-length splice isoforms and APA sites, and higher isoform density than reference genome was identified with SMRT. Recently, SMRT has been used to characterize the complexity of transcriptome in animals and plants. SMRT was applied to analyze human transcriptome and about 14,000 spliced genes were identified, in which over 10% were not previously annotated (9). PacBio SMRT was employed for whole-transcriptome profiling in *Oryctolagus cuniculus* and a total of 36,186 high-confidence transcripts were obtained, among which more than 23% of genic loci and 66% of isoforms have not been annotated before. Furthermore, 24,797 alternative splicing (AS) and 11,184 alternative polyadenylation (APA) events were detected, respectively (10). The first full-length insect transcriptome was sequenced based on the PacBio platform and the first quantitative transcription map of animal mitochondrial genomes was constructed. These transcriptome results enrich fundamental concepts of mitochondrial gene transcription and RNA processing, particularly of the rRNA primary (sequence) structure (11). The sorghum transcriptome was analyzed by the 3^rd^ sequencing technology and results showed that sequencing data uncovered over 7,000 novel alternative splicing events, about 11,000 novel splice isoforms, over 2,100 novel genes and several thousand transcripts that differ in 3’ untranslated regions due to APA (12). Combining with NGS and SMRT, approximately 83.4% of intron-containing genes were found alternatively spliced and the results enhanced understanding on AS under normal condition and in response to ABA treatment in *Arabidopsis thaliana* (8). In moso bamboo, genome-wide AS and APA were identified. The results showed that more than 42,280 distinct splicing isoforms were derived from 128,667 intron-containing full-length non-chimeric (FLNC) reads and 25,069 polyadenylation sites from 11,450 genes, 6,311 of which have APA sites (13). In wheat, a total of 91,881 high-quality FLNC reads were identified and 3,026 new genes were found not annotated previously (14). In maize, 111,151 unique isoforms and higher isoform density than reference genome were identified with SMRT. Moreover, 867 novel high-confidence lncRNAs were identified and had a much longer mean length than those identified by Illumina short-read sequencing (15). Those works provided useful information of transcriptomes and served as valuable resources for further research.

Although work on genome sequencing has been done in red clover, information about sequences and structure of mRNA transcripts is very limited. The genome is not well annotated and number and type of AS events and lncRNAs, the number of splice isoforms, the APA sites and fusion transcripts are largely unclear. Here we used SMRT to analyze red clover transcriptome. Compared with previous annotations, we identified 31,924 novel transcripts, 2,194 of which were derived from novel genes. A total of 5,492 AS events were identified and the majority of AS events was intron retention. We identified 4,333 lncRNAs and the number in reference genome was 11. We also identified 3,762 fusion transcripts, and fusion events were more likely to occur inter-chromosomally. The transcriptome data provided full-length sequences and gene isoforms of transcripts in red clover, which will improve genome annotation and enhance our understanding of the gene structure of red clover.

## Results

### Red clover transcriptome sequencing with SMRT

The transcriptome of 10 pooled samples was sequenced and analyzed with the PacBio Sequel platform to accurately capture full-length sequences and uncover full-length splice variants. RNA from pooled samples isolated and the cDNA was size-selected in fractions of lengths <4-kb and >4-kb. With SMRT, a total of 7,774,277 subreads (13.24-Gb) were obtained, with an average read length of 1,703-bp and N50 of 2,719-bp (Fig. 1a; Table 1). To provide more accurate sequence information, circular consensus sequence (CCS) was generated from reads that pass at least 2 times through the insert, and a total of 525,906 CCS were obtained. By detecting the sequences, 437,814 were identified as full-length (containing 5’ primer, 3’ primer and the poly(A) tail) and 434,405 were identified as full-length non-chimeric (FLNC) reads with low artificial concatemers and the mean length of FLNC was 2688-bp (Fig.1b; Table 1). The FLNC reads with similar sequences are clustered together using ICE (Iterative isoform-clustering) algorithm, and each cluster is considered as a consistent sequence with obtaining 216,028 consensus isoforms (Fig.1c; Table 1). Combined with non-full-length sequences, the quiver program was used to correct the consistent sequences in each cluster, resulting in 17,895 high-quality isoforms (HQs) with accuracy > 99% and 45,883 low-quality isoforms (LQs). Quiver was then used to polish the non-chimeric transcripts and then 215,586 high quality, FL, and polished consensus transcripts were generated. The mean length of polished consensus isoforms was 2657-bp, with N50 of 4616-bp (Table 1). All polished consensus isoforms were corrected using the NGS reads with Proovread software, resulting in 206,465 corrected isoforms, with a N50 length of 4738-bp and mean read length of 2789-bp. In total, 189,960 isoforms (92.0%) were longer than 500-bp, and 155,606 isoforms (75.4%) were longer than 1-kb (Fig.1d; Table 1).

**Figure 1:**
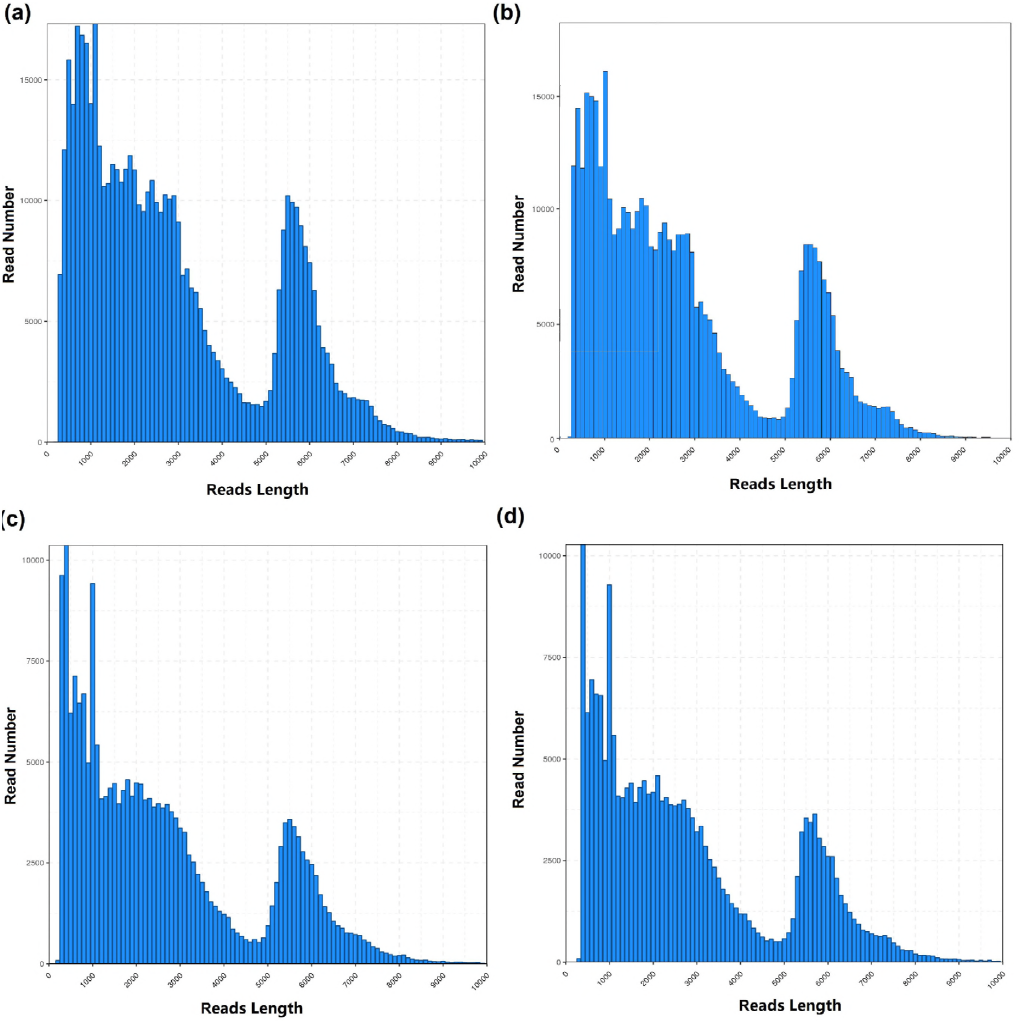
Length distributions of PacBio SMRT sequencing. (**a**) Number and length distributions of 525,906 CCS sequences. (**b**) Number and length distributions of 434,405 FLNC sequences. (**c**) Number and length distributions of 216,028 consensus isoforms. (**d**) Number and length distributions of 206,465 corrected isoforms.

**Table 1:** Summary of ROIs from PacBio single-molecule long-read sequencing.

|  | **Subread** | **CCS** | **FLNC** | **ICE consensus** | **Polished**<br>**consensus** | **Correct**<br>**Consensus** |
| --- | --- | --- | --- | --- | --- | --- |
| **Number** | 7,774,277 | 525,906 | 434,405 | 216,028 | 215,586 | 206,465 |
| **Mean Length** | 1,703 | 2,882 | 2,688 | 2,638 | 2,657 | 2,789 |
| **N50** | 2,719 | 5,026 | 4,840 | 4,591 | 4,616 | 4,738 |

**Table 2:** CDS identification from PacBio single-molecule long-read sequencing.

|  | **CDS** | **Completed CDS** | **3' partial** | **5' partial** | **Uncertain** |
| --- | --- | --- | --- | --- | --- |
| **Number** | 36,622 | 10,304 | 1,929 | 13,158 | 11,231 |

### Functional Annotation of transcripts

All 206,465 transcripts (corrected isoforms) were functional annotated by searching NR, Swissprot, GO, COG, KOG, Pfam and KEGG databases and a total of 199,093 transcripts (96.4%) was annotated (Fig. 2a; Table S1). We analyzed homologous species by comparing the transcript sequences to the NR database, and the results showed that the largest five number of transcripts was distributed in *Trifolium subterraneum* (84,418), *Medicago truncatula* (57,646), *Cicer arietinum* (17,825), *Trifolium pratense* (10,284) and *Glycine max* (1,726) (Fig. 2b). GO analysis showed that the enrichment of 80,667 transcripts could be divided into three groups, including biological processes, molecular functions and cellular components. Genes involved in biological processes consisted of metabolic, cellular, single-organism, localization, biological regulation, signal processes and so on (Fig. 2c). Genes in the molecular function were mainly enriched for binding, catalytic, transporter, structural molecular and nucleic acid binding transcription factor activities. For the category “Cellular Component”, genes were mainly involved in cell, cell part, membrane, organelle, membrane part and others. The KEGG results demonstrated that 186,497 transcripts were mapped to 368 KEGG pathways (Fig. 2d).

**Figure 2:**
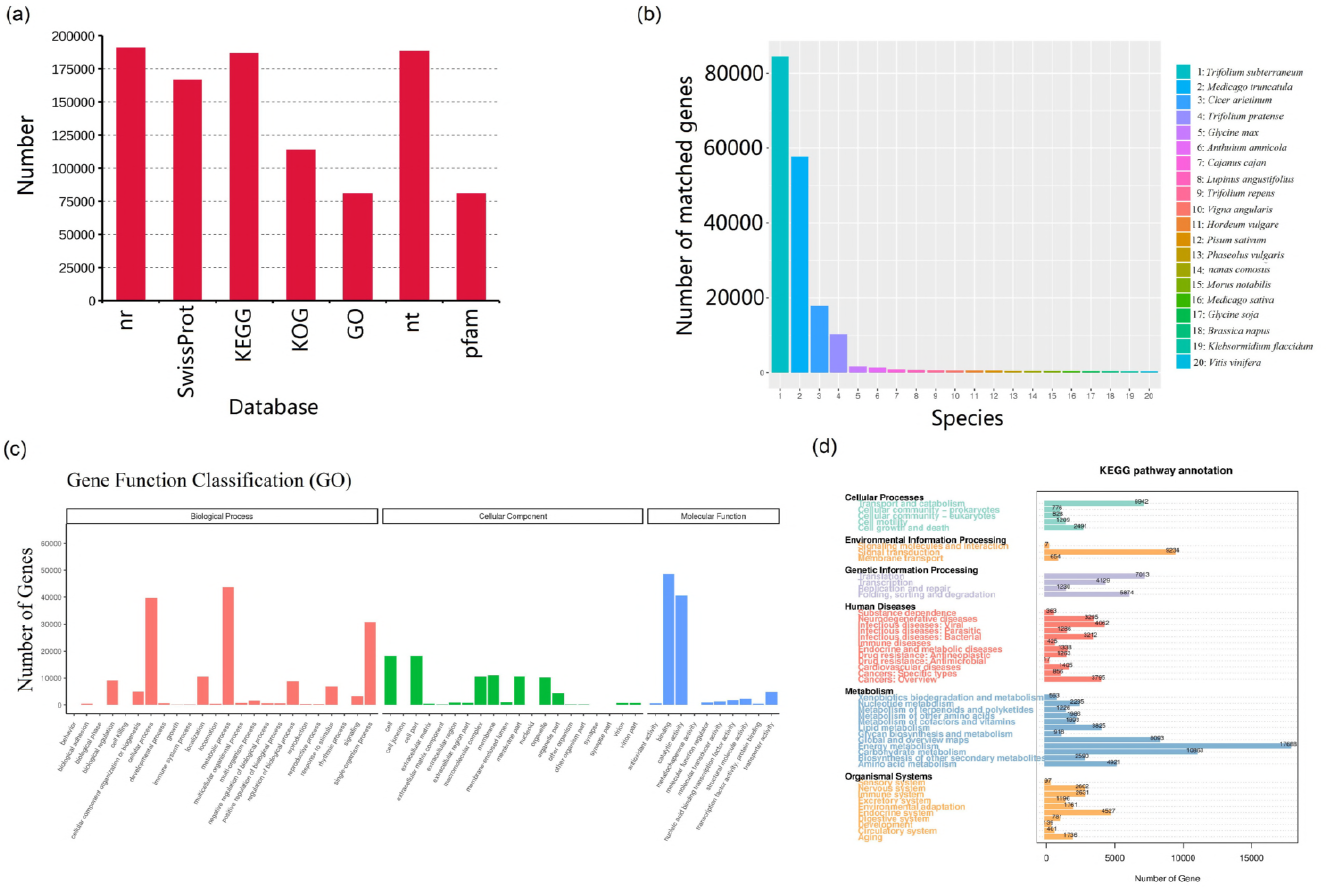
Function annotation of corrected isoforms. (**a**) Function annotation of transcripts in all databases. Nr, Non-Redundant Protein Database; GO, Gene Ontology; COG/KOG, Cluster of Orthologous Groups of proteins; KEGG, Kyoto Encyclopedia of Genes and Genomes. (**b**) Nr Homologous species distribution diagram of transcripts. (**c**) Distribution of GO terms for all annotated transcripts in biological process, cellular component and molecular function. (**d**) KEGG pathways enriched of transcripts.

### Genome Mapping

We compared all 206,465 corrected isoforms against the draft genome sequence of red clover using GMAP. A total of 136,395 reads (66.06%) were mapped to the reference genome. Based on the mapped results, these reads could be divided into four groups (Fig. 3a). Unmapped, consisted of 70,070 reads (33.94%) with no significant mapping to the draft genome. Multiple mapped, contained 130,525 reads (63.22%) showing multiple alignments. Mapped to ‘+’, included 4,414 reads were mapped to the positive strand of the genome and mapped to ‘-’, included 1,456 reads were mapped to the opposite strand of the genome. Isoforms spanning two or more genes are removed from downstream splice isoform analysis; however, they likely represent misannotations in the gene models. In this case, 48,626 isoforms were removed, and the remaining 87,769 transcripts were then assembled into clusters, resulting in 39,787 isoforms. By analysis, 89.2% of those reads aligned to 13,284 annotated genes out of 40,868 genes in the reference genome.

**Figure 3:**
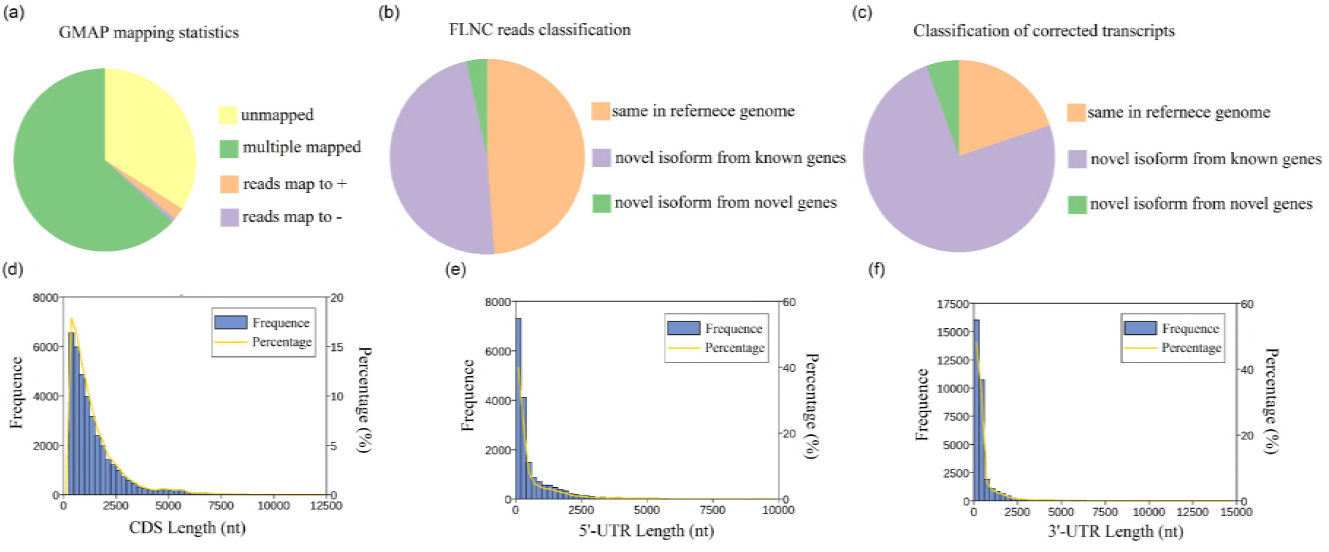
GMAP analysis and CDS-UTR structures of SMRT sequences. (**a**) GMAP analysis of corrected reads to reference genome. (**b**) Classification of FLNC sequences mapped to reference genome. (**c**) Classification of corrected isoforms mapped to reference genome. (**d**) Number, percentage and length distributions of coding sequences of corrected isoforms. (**e**) Number, percentage and length distributions of 5’-UTR of corrected isoforms. (**f**) Number, percentage and length distributions of 3’-UTR of corrected isoforms.

### Novel genes and transcripts finding

All FLNC sequences were compared against the genome sequence with GMAP, and 200,787 reads (46.22%) were mapped to the reference genome. Mapped FLNC were divided into 3 types: 1) 98,030 same in reference genome; 2) 95,898 novel isoforms from known genes; 3) 6,859 novel isoforms from novel genes (Fig.3b). We also compared 39,787 isoforms against the reference genome, and 89.2% of those reads aligned to 13,284 annotated genes out of 40,868 genes in the reference genome. We identified 7,863 same in reference genome, 29,730 novel isoforms from known genes and 2194 novel isoforms from novel genes (Fig.3c; Fig.4).

### CDS finding and Exon/Intron structure analysis

Coding sequences were identified by ANGEL software, resulting in 36,622 coding sequences with a mean length of 1435.26 nucleotides (Fig.3d). Sequences with start and stop codons were defined as complete coding sequences, and 10,304 carried complete ORFs (Table 2). The number and length distributions of 5’ and 3’ UTRs were investigated. The results displayed 18,198 5’ UTRs with a mean length of 650.5-bp and 33,219 3’ UTRs with a mean length of 559.45-bp (Fig.3e-f).

By SMRT, 39,787 transcripts were used to analyze exon structure. One exon was found in 8,302 transcripts (20.87%) and the transcript number with 2 exons was 5,175 (13.01%), while transcripts with more than 20 exons was 1,927 (4.84%) (Fig. 5a; Table S2). In reference genome, the transcripts number with 1 exon was 40,868 (49.18%) and transcripts with more than 20 exons was 7,693 (9.26%) (Fig. 5b). The intron number was 596,960 in reference genome with 14.61 introns per gene (intron/gene) and that was 247,837 in our Iso-Seq data with 16.27 intron/gene. We further analyzed the median number of introns in intron containing genes, and the result showed the number is five in red clover, which is four in *Arabidopsis thaliana*.

**Figure 5:**
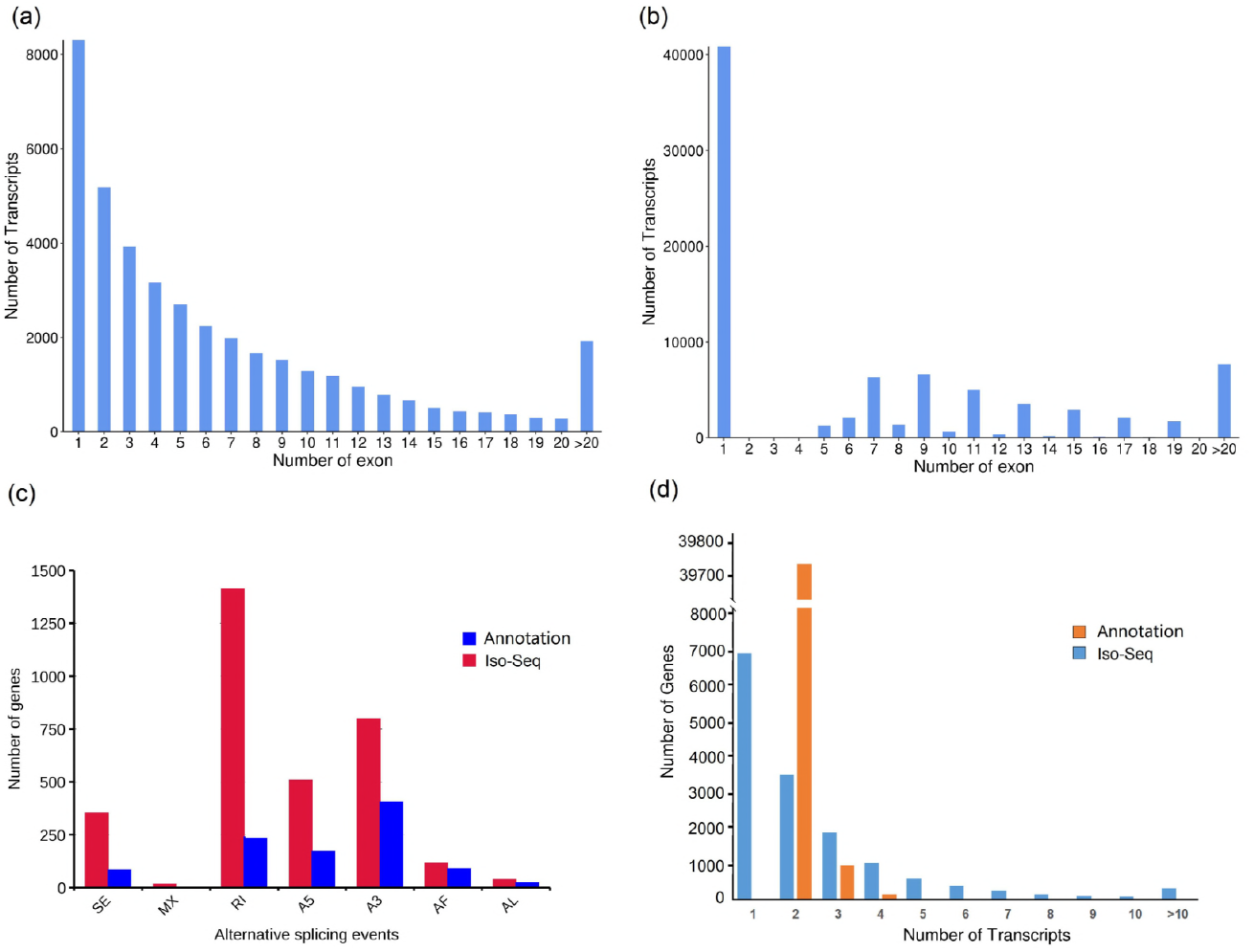
Gene structures and AS events. (**a**) Distribution of transcripts that have one or more exons from Iso-Seq. (**b**) Distribution of transcripts that have one or more exons from reference genome. (**c**) The total number of AS events in detected genes by SMRT compared with the annotated gene models. Annotation, AS events in genes detected by SMRT based on reference genome; Iso-seq, AS events in genes detected by SMRT based on Iso-Seq reads. SE, skipped exon; MX, mutually exclusive exon; A5, alternative 5’ splice site; A3, alternative 3’ splice site; RI, retained intron; AF, alternative first exon; AL, alternative last exon. (**d**) Distribution of genes that produce one or more splice isoforms from Iso-Seq and annotation gene models.

### AS and Splice isoforms in red clover

AS events in red clover were analyzed with Suppa software. We detected a total 5,492 AS events from the Iso-Seq reads and there are 1,123 AS events in reference genome. The majority of AS events in Iso-Seq was intron retention with similar distribution to other plants, while the majority of AS events in reference genome was alternative 3’ splice (Fig.4; Fig. 5c). Compared with reference genome, 4,831 AS events occurred specifically in 1,996 genes based on Iso-Seq data (Table S3). Those AS events in our study largely enriched transcripts information in the draft version of the red clover genome. In reference genome, there are 40,868 genes annotated and 39,729 genes were shown to have 2 isoforms. The largest number of isoforms was 6 found in 3 genes, Tp57577_TGAC_v2_gene16492 (Tp57577_TGAC_v2_LG7: 11814616-11818691), Tp57577_TGAC_v2_gene3878 (Tp57577_TGAC_v2_LG6: 17447258-17473786) and Tp57577_TGAC_v2_gene39688 (Tp57577_TGAC_v2_scaf_1548: 11427-14327). In our Iso-Seq analysis, only one single isoform was detected in 6,904 genes and two or more isoforms were found in 8,325 genes. Six and more than 6 splice isoforms were detected in 1,329 genes (Fig. 5d; Table S4). For example, six isoforms were detected in Tp57577_TGAC_v2_gene25740 based on Iso-Seq, while the isoform number was 3 in the red clover annotation (Fig. 5e). Isoforms per gene (2.61) identified from Iso-Seq data were significantly more than that in the reference genome. The largest number of isoforms were 113 found in Tp57577_TGAC_v2_gene14962 (Tp57577_TGAC_v2_scaf_591:48683-68918).

### Identification of lncRNA, fusion transcript and TF

A total of 4,333 lncRNAs were identified by three methods and 3,050 (70.39%) of the lncRNAs were single exon (Fig. 6a-c; Table S5). We classified them into 4 groups: 1,148 lincRNA (26.49%), 289 antisense (6.67%), 1,747 sense intronic (40.32%) and 1,149 sense overlapping lncRNAs (26.52%) (Fig. 6b). BLASTN was used to get rid of the previously discovered 11 lncRNAs downloaded from ensemble website, and all predicted lncRNA (4,333) were identified as novel lncRNAs with a mean length of 665.39-bp (Fig. 6c-d). Mapping lncRNAs to chromosomes was shown in Fig. 4, and identification of lncRNAs enriched genome information of red clover.

**Figure 4:**
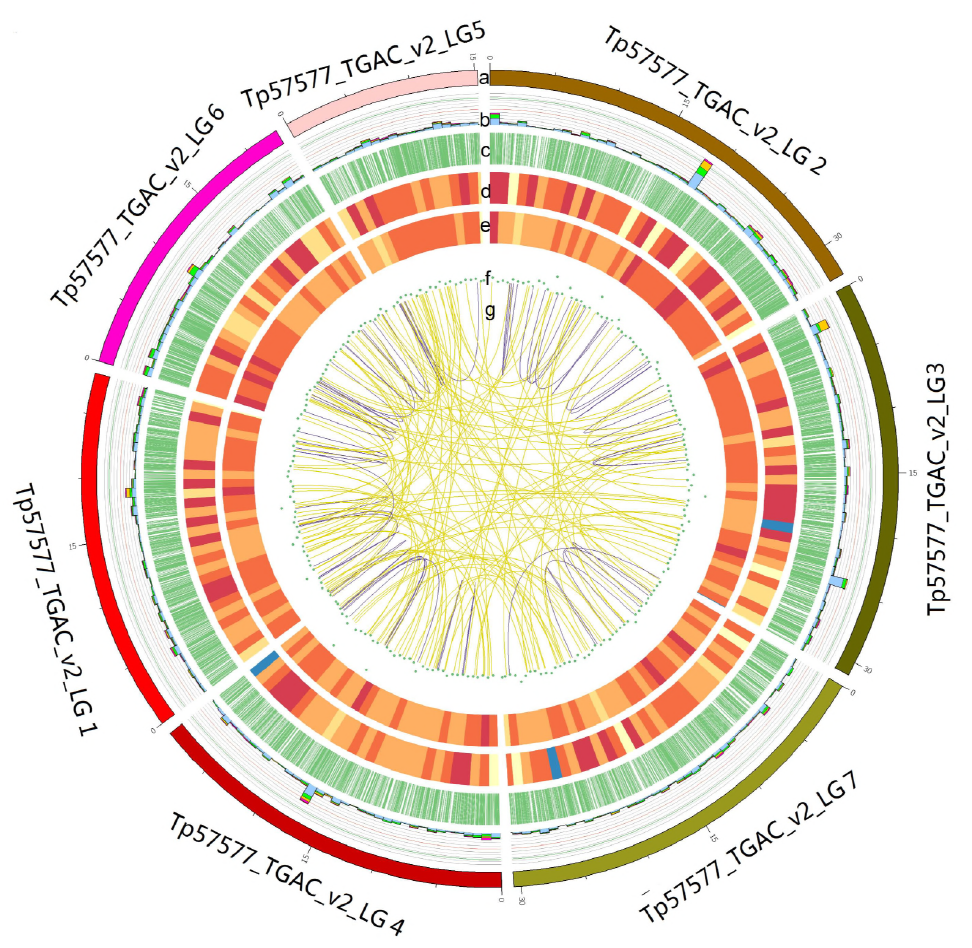
CIRCOS visualization of Iso-Seq data at the genome-wide level. (**a**) Karyotype of red clover chromosomes. (**b**) Comparison of APA sites mapped to red clover reference genome. (**c**) Novel transcript density from Iso-Seq data. The closer to red, the higher density the color represents and the closer to blue, the lower density the color represents. (**d**) Novel gene density from Iso-Seq data. (**e**) LncRNA density. The closer to the center, the lower density the dot represents. (**f**) Linkage of fusion transcripts. Purple, intra-chromosomal; Yellow, inter-chromosomal.

**Figure 6:**
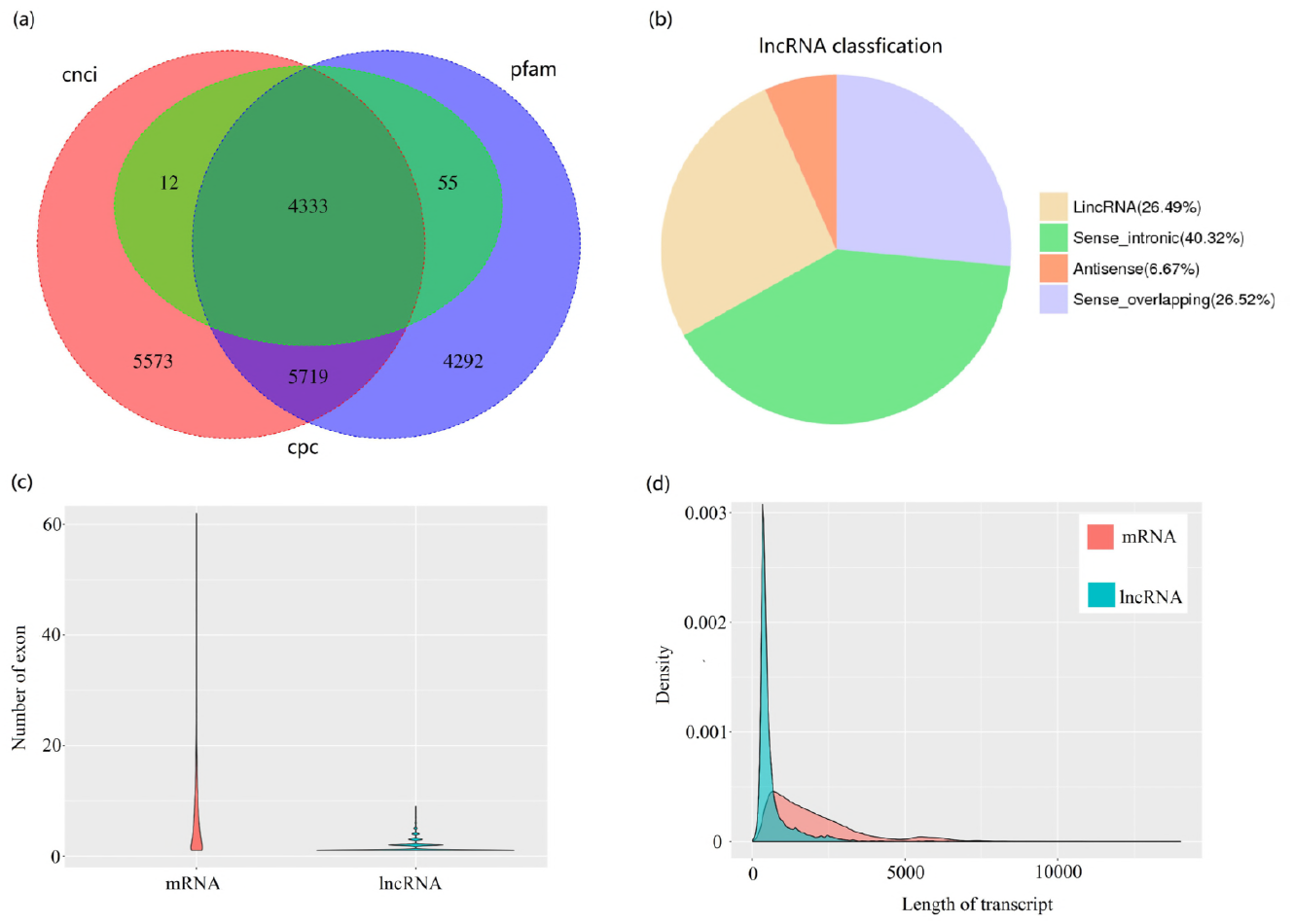
Identification of lncRNAs. (**a**) Venn diagram of lncRNAs predicted by CPC, CNCI and pfam methods. (**b**) Proportions of four types of lncRNA. (**c**) Comparison of exon number of mRNA and lncRNAs identified in this study. (**d**) Density and length distributions of mRNAs and lncRNAs in red clover.

In reference genome, a total of 39,051 scaffold sequences was used to predict fusion transcript. In this study, we identified 3,762 fusion transcripts out of 39,051 scaffold sequences, and fusion events were more likely to occur inter-chromosomally (3,665) than intra-chromosomally (97). Seven scaffolds, assembled to chromosome level from 39,051 scaffold sequences, were picked up for fusion transcript identification. We also found 334 fusion transcripts, including 238 inter-chromosomal and 96 intra-chromosomal sequences (Fig. 4). To further validate the fusion transcripts, we randomly chose 10 candidates and primers were designed based on the sequences of Iso-Seq. Eight (80%) were validated by RT-PCR and Sanger sequencing (Fig. S1). For example, i0_HQ_c62675/f4p0/1006 was fused by two gene TP57577_TGAC_V2_GENE20501 and TP57577_TGAC_V2_GENE12817 (Table S6). The PCR products were confirmed by Sanger sequencing and the results proved the authenticity of these chimeric RNAs.

Transcript factors (TFs) and transcript regulators (TRs) play important regulatory roles in plant growth and development. They were identified and classified with iTAK and 2,302 putative TF (1,361) and TR (941) members from 89 families were identified (Table S7). The top 29 families identified were shown in Fig. 7a. We compared those members against 2,065 known TFs from red clover, and the result showed that 249 were identified as known and 2,053 were identified as novel.

**Figure 7:**
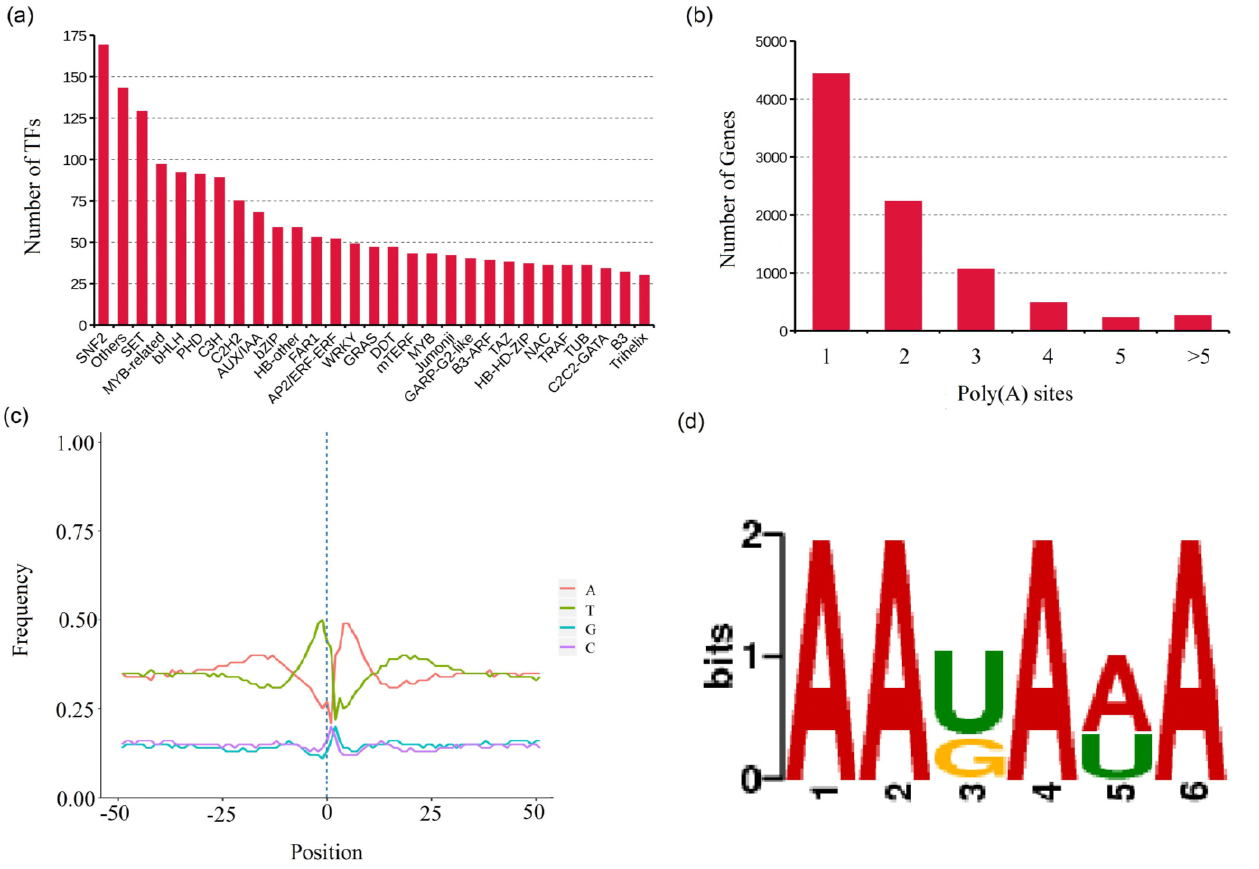
TF prediction and APA analysis. (**a**) Number and family of top 29 TFs predicted by SMRT. (**b**) Distribution of the number of poly(A) sites per gene. (**c**) Nucleotide composition around poly(A) cleavage sites. The relative frequency of a nucleotide is shown as a function of genomic position across all poly(A) cleavage sites detected in our data. (**d**) MEME analysis of an over-represented motif at 25-nts upstream of the poly(A) site in red clover transcripts.

### APA analysis

To identify accurately differential polyadenylation sites in red clover, 3’ ends of transcripts from Iso-Seq were investigated. Of the 15,229 genes detected by Iso-Seq, 8,719 including 1,86,517 transcripts have at least one poly(A) site, and 497 genes have at least five poly(A) sites (Fig. 4; Fig. 7b; Table S8). The average number of poly(A) site per gene was 1.97. We analyzed the nucleotide composition in the upstream (50 nts) and downstream (50 nts) of all poly(A) cleavage sites for nucleotide bias. Nucleotide composition flanking all poly(A) sites was analyzed. Consistent with findings in *Sorghum bicolor* and *Phyllostachys edulis*, we observed clear nucleotide bias around poly(A) sites in red clover with an enrichment of uracil (U) upstream and adenine (A) downstream of the cleavage site in 3’-UTRs (Fig. 7c). To identify potential cis-elements necessary for polyadenylation, we performed a MEME analysis for motifs enriched upstream of the cleavage site using 50 nucleotides upstream from the predominant poly(A) site of all transcripts. One conserved motif (AAUAAA) was identified upstream of poly(A) cleavage sites, and this motif was consistent with previous reported patterns in *Sorghum bicolor*, and *Phyllostachys edulis* (Fig. 7d).

## Discussion

By now, the draft genome sequence had been published, but the transcriptome was not fully explored. The full-length mRNA sequences, alterative spliced transcripts, APA sites and fusion transcripts in red clover remains unknown. With development of sequencing technology, single molecule long reads sequencing technology provides new insights into full-length sequence, alterative splice, gene structure and APA. In this work, analyzed full-length transcriptome of red clover with PacBio SMRT. PacBio sequencing yielded 525,906 CCS, in which 434,405 are identified as FLNC transcripts. The length of the FLNC sequence reflects the length of the cDNA sequence in each sequencing library, and the library can be assessed by the length of the FLNC sequence. The length of the FLNC sequence is consistent with the size of the library (Fig.1). A total of 216,028 consensus isoforms were generated with ICE algorithm and 215,586 polished consensus were identified.

Combined SMRT with NGS, 206,465 corrected consensus reads were obtained in total. Meanwhile, 5,492 AS events, 4,333 lncRNAs, and 3,762 fusion transcripts were identified and 8,719 genes with 1,86,517 transcripts have at least one poly(A) site. Those new findings provided important information for improving red clover draft genome annotation and fully characterization of red clover transcriptome.

Full-length transcript sequence information is very useful for both genome annotation and gene function studies in plants. Our results demonstrated that PacBio sequencing is an effective technology for obtaining reliable full-length transcript sequence information in red clover. PacBio sequencing yielded 525,906 CCS, 82.6% of which are identified as FLNC transcripts. PacBio CCSs and FLNC reads completely avoided the need to assemble short NGS reads. With NGS data, 206,465 corrected isoforms were obtained, and 75.4% were longer than 1-kb. About 10,304 isoforms were found to carry completed ORF. The high capacity of PacBio transcriptome sequencing to generate full-length transcript sequence information may well be related to its long-read property. In our work, 15,229 genes were detected by SMRT with a mean of 3,788-bp, which is 405-bp larger in size than that in reference genome. Previous reports supplied similar suggestion, showing that newly discovered transcripts by SMRT in wheat were on average more than 45-bp longer than the known transcripts (14). The new sequencing technology enriches transcript resources and provides advantages for discovering novel or uncharacterized transcript isoforms and genes. In this work, we identified 29,730 novel isoforms from known genes and 2194 novel isoforms from novel genes in the draft genome based on SMRT data. Those findings not only enrich the transcriptional information of the draft genome sequence but also are useful for functional studies of important genes in further research.

Previous studies on red clover transcriptome mainly focused on NGS technology to exploring gene discovery, differential expression genes analysis, pathways enrichment, marker identification and so on (6, 7). But this tool is often unable to accurately capture or assemble full-length transcripts. Based on PacBio SMRT, fragmenting of RNA is not required and intact transcript sequence information is provided by avoiding assembly. In our work, we characterized red clover transcriptome with PacBio SMRT. The N50 and average lengths of contigs assembled from previous project were 1707 and 1262-bp (7), and those were much larger from Iso-seq (Table 1). Those results showed that SMRT has a better capacity in capturing transcript sequences, especially long transcript sequences. As well as full-length transcripts, the AS event identification is another advantage. We detected a total 5,492 AS events from the Iso-Seq reads and there are 1,123 AS events in reference genome. The majority of AS events in Iso-Seq is intron retention with similar distribution to maize (15), bamboo (13), *Amborella trichopoda* (16), strawberry (17) and so on, while the majority of AS events in reference genome is alternative 3’ splice. Those AS events in our study largely enriched transcripts information in the draft version of the red clover genome. But we did not analyze the isoform expression levels in current project, so detailed expression levels of isoforms derived from one gene in red clover should be analyzed combined expression analysis from NGS with AS isoforms from SMRT in the future.

In our study, we found 3,762 fusion transcripts, and fusion events were more likely to occur inter-chromosomally than intra-chromosomally. The chimeric fusion events in red clover enhanced the complexity of the red clover transcriptome and those results were consistent with the higher proportion of inter-chromosomal to intra-chromosomal fusions in maize (15). To confirm the fusion transcripts, we randomly selected 10 candidates to design primers for RT-PCR analysis. As predicted, 8 candidates were validated by RT-PCR and Sanger sequencing. The findings in fusion transcript, would greatly improves draft version of gene models in red clover. LncRNAs, a hotpot of molecular biology, is thought to be important regulators with function little known. In this study, we identified 4,333 lncRNAs with a mean length of 665.39-bp and the largest type in number is sense intronic. We downloaded 11 known lncRNA from ensemble website, the mean length is 93.09-bp. By comparison, the newly identified lncRNAs were much longer, with a mean length 7-time larger than that of known lncRNAs. Those results is similar to the former study. In maize, lncRNAs identified by SMRT, most of which are intergenic, are much longer than those previously described, and some of which are as long as 6-kb (15). Of the 15,229 genes detected by Iso-Seq, 8,719 including 1,86,517 transcripts have at least one poly(A) site (Fig.4; Fig. 7b). This work provides useful information for future analysis of the relation of APA and gene function.

In conclusion, we analyzed full-length transcriptome of red clover with PacBio SMRT. Those new findings provided important information for improving red clover draft genome annotation and fully characterization of red clover transcriptome.

## Materials and Methods

### Plant samples

Red clover seeds (cv. Common) received as gifts from Dr. Feifei Li of Top Green Group (Beijing, China) were sown in nutrition medium containing peat, vermiculite and pearlite (1:1:1) until germination, and then plants were grown at Beijing Forestry University (E116°20’; N40°00’) under greenhouse conditions at 25/23°C (day/night) with a 16-h photoperiod in growth chambers. Different tissues, including leaves, stems, roots and flowers of 1-year-old plants were harvested in May 2017. To obtain more spliced isoforms, red clover plants were stressed by 100mM NaCl or 10% PEG for 1 day, or induced by 10μM ABA, 10μM GA_3_, 10μM IAA and 10μM 6-BA for 6 hours. All samples were harvested and frozen in liquid nitrogen and stored at −80°C for further experiments.

### Library preparation and SMRT sequencing

Total RNA from each sample was isolated using a Plant RNA kit (Omega bio-Tech, USA) and then treated with RNase-free DNase I (NEB) to remove contaminated genomic DNA. RNA degradation and contamination was monitored on 1% agarose gels and RNA purity was checked using the NanoPhotometer^®^ spectrophotometer (IMPLEN, CA, USA). RNA concentration was measured using Qubit^®^ RNA Assay Kit in Qubit^®^ 2.0 Flurometer (Life Technologies, CA, USA) and RNA integrity was assessed using the RNA Nano 6000 Assay Kit of the Bioanalyzer 2100 system (Agilent Technologies, CA, USA). A total amount of 5 μg of total RNA (equally mixed with all RNAs) was used as input into the Clontech SMARTer reaction. Size fractionation and selection (<4kb and >4kb) were performed using the BluePippin™ Size Selection System (Sage Science, Beverly, MA). Two SMRT bell libraries were constructed with the Pacific Biosciences DNA Template Prep Kit 2.0 and SMRT sequencing was then performed on the Pacific Bioscience Sequel System.

### Quality filtering and Error Correction

Raw reads was processed using the SMRTlink (version 4.0) software with parameters: minLength=200, minReadScore=0.75. CCSs were generated from subread BAM files (parameters: min_length 200, max_drop_fraction 0.8, no_polish TRUE, min_zscore −999, min_passes 1, min_predicted_accuracy 0.8, max_length 18000) and a CCS. BAM file was output. By identifying the 5’ and 3’ adapters and the poly(A) tail, CCS were then classified into full length and non-full length reads. A full-length read contained both the 5’ and 3’ primers and there was a poly(A) tail signal preceding the 3’ primer. CCS with all three elements and not containing any additional copies of the adapter sequence within the DNA fragment are classified as FLNC. Then, consensus isoforms were identified using the algorithm of ICE (Iterative Clustering for Error Correction) from FLNC and were further polished with non-full length reads to obtain high-quality isoforms with post-correction accuracy above 99% using Quiver (parameters: hq_quiver_min_accuracy 0.99, bin_by_primer false, bin_size_kb 1, qv_trim_5p 100, qv_trim_3p 30). The Illumina RNA-Seq data (SRA submission number:

SUB2425623) generated by our lab was used to correct nucleotide indels and mismatches in consensus reads with the software Proovread (Version 2.12) (18), resulting in corrected isoforms.

### Mapping to the reference genome and gene structure analysis

FLNC and corrected isoforms were aligned to the reference genome with the Genome Mapping and Alignment Program (GMAP, version: 2017-01-14) (19) with parameters: --no-chimeras, --cross-species, --expand-offsets 1-B 5 -K 50000 -f samse -n 1. The output files are in the BAM format. Gene structure analysis was performed using TAPIS pipeline (Version 1.2.1, <u>https://bitbucket.org/comp_bio/tapis</u>) (12). The gff3 format genome annotation file was transfer into GTF format with gffread (20) and then used for gene and transcript determination. Alternative splicing events were identified using the SUPPAA (Version: 2017-02-07) (21). SUPPA generates different alternative splicing event types: exon skipping (SE), alternative 5’ and 3’ splice sites (A5/A3), mutually exclusive exons (MX), intron retention (RI), and alternative first and last exons (AF/AL). The exon-intron structure for each transcript were predicted and the numbers of introns were statistically analyzed in a transcriptome level. APA events were then analyzed by TAPIS described previously (12). To identify poly(A) sequence signals in Iso-Seq data, MEME analysis was performed on the sequence of 50 nucleotides upstream of the 1,86,517 poly(A) sites. MEME-ChIP was run locally on a Linux system described previously (12). Fusion transcripts were determined as transcripts mapping to two or more long-distance range genes and was validated by at least two Illumina reads described as previously (15).

### Functional Annotation

Corrected isoforms were searched against NCBI non-redundant (NR), NCBI nucleotide sequence (NT),

Swiss-Prot, Cluster of Orthologous Groups (KOG/COG)(22) and Kyoto Encyclopedia of Genes and Genomes (KEGG, version 58) (23) databases with BLAST software (version 2.2.26) under a threshold E-value ≤10-5. Gene Ontology (GO) annotations were determined based on the best BLASTX hit from the NR database using the Blast2GO software (version 2.3.5, E-value ≤10-5) (24). KEGG pathway analyses were performed using the KEGG Automatic Annotation Server (KAAS1) and HMMER software (25) was used to search Pfam database.

### Identification of ORFs, TFs and lncRNAs

Corrected isoforms were used for further analysis. To predict ORFs, the ANGEL pipeline, a long read implementation of ANGLE, was used to find potential coding sequences from transcripts (26). Those transcripts containing complete ORFs as well as 5’- and 3’-UTRs (untranslated regions) were designated as full-length transcripts and the putative protein sequences were predicted. The transcription factors were predicted with iTAK software (27) from putative protein sequences. A total of 2065 known red clover TFs were downloaded from PlantTFDB (Plant Transcription Factor Database v4.0) (28), and blastp was ran with the following cutoff criteria: E-value ≤10-5, min-coverage=85% and min-identity= 90%. The red clover lncRNAs were downloaded from ensemble website (<u>ftp://ftp.ensemblgenomes.org/pub/plants/release-37/fasta/trifolium_pratense/ncrna/</u>) and all 11 known lncRNAs were identified as sense intronic type. To identify long non-coding RNA (lncRNA) in the PacBio data, three analysis methods including CPC (29), CNCI (30) and Pfam (31) were used and then transcripts with length less than 300-bp predicted in all 3 methods were removed. BLASTN was used to get rid of the previously discovered lncRNAs under a criteria of e-value ≤1e-10, min-identity=90% and min-coverage=85%. The lncRNAs were divided into four groups: lincRNA, antisense, senese intronic and sense overlapping based on the method reported by Mathew (32).

### RT-PCR validation of fusion transcripts

For PCR validation of fusion transcripts, gene-specific primers were designed using DNA Man (Version 6.0) and Primer Premier (Version 5.0). All primers used in the RT-PCR analyses are reported in Supplementary Table S9.

## Acknowledgment

The program was supported by the National Natural Science Foundation of China (No. 31601989 and No. 31672477).

## Author contribution statement

L. H. and L. X. conceived and designed the research. Y. C. and J. Y. conducted experiments. S. L. and S. J. analyzed data. Y. C. wrote the manuscript. All authors read and approved the manuscript.

## Conflict of interest statement

The authors declare that there is no conflict of interest.

## Abbreviations

ORF: open reading frames
lncRNA: long non-coding RNAs
TF: transcript factor
NGS: next-generation high-throughput sequencing
SMRT: single molecule long reads sequencing technology
AS: alterative splice
APA: alternative polyadenylation
ABA: abscisic acid
FLNC: full-length non-chimeric
CCS: circular consensus sequence
TF: Transcript factor
TR: transcript regulator.

